# Targeting Estrogen to the Brain via the Prodrug DHED does not Protect Against Metabolic Dysfunction in Obese, OVX mice

**DOI:** 10.64898/2026.01.26.701850

**Authors:** Celine Camon, Elodie Kip, Rebecca Lord, Caroline Decourt, Mel Prescott, Jenny Clarkson, Katalin Prokai-Tatrai, Stephanie M. Correa, Rebecca E. Campbell, Michael Garratt

## Abstract

Menopausal hormone therapy (MHT) is prescribed for climacteric symptoms including hot flushes and weight gain and contains estrogens such as 17 beta-estradiol (17βE2). However, estrogen receptor activation by MHT may increase reproductive cancers and cardiovascular event risk in some people. As the protective metabolic effects of 17βE2 are partly mediated through the arcuate nucleus of the hypothalamus, restricting 17βE2 actions to the brain could serve as a safer mechanism of MHT. 10β,17β-Dihydroxyestra-1,4-dien-3-one (DHED) is a prodrug of 17βE2 which is enzymatically converted to the parent hormone exclusively within the brain. DHED has demonstrated positive benefit in rodent models of centrally-mediated maladies including hot flushes, depression and cognitive decline, without peripheral hormonal burden. Therefore, we hypothesized that DHED treatment in obese female mice would act within the hypothalamus to provide the same beneficial metabolic effects as 17βE2. Female mice were ovariectomized, placed on a high fat diet and split into either control, DHED, or 17βE2 treatment groups. Body weight, uterus weight and glucose tolerance were recorded along with gonadal hormone receptor expression in the brain. Delivery of DHED at a similar dose as 17βE2 failed to improve metabolic parameters or recapitulate the hypothalamic responses induced by 17βE2. Delivery of DHED at higher doses, which elicited estrogen-like actions within the brain, still failed to improve metabolic health. Our findings suggest that peripheral actions, in addition to hypothalamic targets, may be required to mediate 17βE2’s protective effects on metabolism and that brain-targeted MHT may be unsuitable for improving metabolic health during menopause.

## Introduction

The reduction of circulating ovarian hormones, loss of follicular reserve and the eventual cessation of menstruation mark the transition to menopause ^1^. Hormonal changes that occur over this period include the relatively abrupt decline of estrogen and progesterone, with the simultaneous reduction in inhibin and anti-Müllerian hormone and an increase in follicle-stimulating hormone ^2^. Many people experience symptoms that can significantly alter their quality of life during this transition, including hot flushes, significant weight gain, sleep disturbances and mood alterations, which are thought to be driven (either directly or indirectly) by estrogen deprivation to the brain ^3,4^.

A shift occurs in metabolic function during menopause, which leads to adipose tissue redistribution from femoral and gluteal subcutaneous deposits and results in increased visceral adiposity ^5^. Other markers of poor metabolic health are also observed during and after menopause, such as elevated cholesterol, impaired glucose tolerance and increased inflammatory markers ^6,7^. These alterations can increase insulin resistance, lipolysis and elevate the overall risk for metabolic disease ^8,9^. The dramatic decline of ovarian 17 beta-estradiol (17βE2) production is thought to play a significant role in the redistribution of adiposity and associated metabolic disturbances during this period^10^. This is mirrored in ovariectomized (OVX) rodents that demonstrate increased abdominal fat, increased body weight and adiposity, along with impaired glucose tolerance ^11–15^. When OVX mice on a high fat diet (HFD) are treated with 17βE2, they show significantly reduced body weight, adipocyte size, food intake, hepatic triglyceride levels and a marked improvement in glucose tolerance, highlighting the therapeutic potential of estrogen replacement in managing metabolic health in menopause ^16,17^. While the safety profile of menopausal hormone therapy (MHT) has been well characterized over the last decade, MHT can be unsuitable for people with, or histories of, reproductive cancers ^18^. Increased cancer risk stems from undesirable hormone-receptor signalling actions within estrogen-sensitive tissues in the periphery, including the uterus and breast^19^. Therefore, alternatives to traditional hormonal treatments are required to provide safe and effective treatments to alleviate menopausal symptoms.

The specific tissues where 17βE2 acts to improve metabolic health are not fully established. However, the hypothalamus, and the arcuate nucleus (ARC) in particular, is likely to play a major role due to its importance in the regulation of body weight, metabolism and food intake ^20 21,22^. Estrogen receptor alpha (ERα), the canonical estrogen receptor activated by 17βE2, is highly expressed in the ARC ^23–25^. Neuron specific knockdown of ERα induces abdominal obesity and hyperphagia in female mice ^26^, and ERα deletion specifically from anorexigenic pro-opiomelanocortin *(POMC)* neurons also induces hyperphagia ^26^. Neurons in the ARC are responsible for sensing circulating levels of leptin, insulin and glucose ^27^. ERα-expressing neurons in the neighboring ventromedial hypothalamus (VMH) of the hypothalamus have also been shown to sense glucose levels and prevent hypoglycemia in female mice, along with playing a key role in thermoregulation ^28,29^. Given that the VMH projects to the ARC, it is likely that metabolic function is also modulated at least partially by this brain region^30^.

10B, 17B-dihydroxyestra-1,4-dien-3-one (DHED) is a unique prodrug of 17βE2 that selectively produces the hormone in the brain to treat neurological conditions in rodent models of hot flush, stroke, cognitive decline and depression ^31–34^. Consequently, administration of DHED does not increase uterine weight or activate ERα in peripheral tissues when assessed using reporter mice with luciferase expression that is dependent on estrogen response element (ERE) activation ^31,34^. The conversion of DHED to 17βE2 occurs via a CNS-specific enzyme, a short-chain NADPH-dependent dehydrogenase/reductase that is expressed exclusively within and abundantly across the brain, including within the hypothalamus ^31^. Therefore, DHED may be a novel method of delivering 17βE2 to hypothalamic centres involved in mediating metabolic health, without acting on peripheral tissues.

We hypothesized that DHED treatment would recapitulate the protective metabolic effects of 17βE2 in obese OVX female mice, while bypassing the activation of peripheral estrogen-sensitive organs. We assessed whether DHED treatment could improve a range of metabolic health parameters including body weight, food intake, glucose tolerance and liver adiposity in OVX mice on a HFD, and how these responses compared to those who underwent 17βE2 treatment. ERα-dependent central effects were assessed by the presence of progesterone receptor (PR) expression, as the expression of PR is largely dependent on estradiol signaling ^32,35^.

Previous proof-of-concept studies investigating the utility of DHED-derived 17βE2 to provide protection against cognitive decline, hot flushes and depression have used equivalent doses of DHED to 17βE2 ^31,32^. We therefore first tested whether DHED treatment at the same dose as 17βE2 would recapitulate estrogen-like actions within the hypothalamus and on metabolic health. We then performed an acute 48-hour dose-response study to identify a dose with sufficient activation within the brain. We subsequently performed another chronic study with increasing doses of DHED on a HFD to investigate whether optimized dosages of this brain selective prodrug of the hormone could provide protection against metabolic dysfunction.

## Methods

### Animal husbandry

All research was approved by the University of Otago Animal Ethics Committee (AUP-22-42) (Experiments 1 and 3) or by the UCLA Institutional Animal Care and Use Committee (IACUC) (Experiment 2). All animals were of the C57BL/6J strain, sourced from either the Biomedical Research Facility (University of Otago, experiments 1 and 3), or bred in-house or sourced directly from Jackson Laboratories (UCLA, experiment 2). Mice were housed in a temperature (21°C) and light-controlled environment on a 12:12 light: dark cycle and had *ad libitum* access to food and water.

### Experimental timelines

#### Experiment 1: Metabolic function and hypothalamic response with a matched dosage of DHED and 17βE2

Female C57BL/6J female mice were ovariectomised (OVX, n=20) between 6-8 weeks of age (**Fig 1a**). Prior to OVX, mice were fed a normal chow diet (Teklad Global 18% Protein Rodent Diet #2918 (Protein: 18.6%; Fat 6.2%; Carbohydrate: 44.2%). Mice were then placed on a 45% high fat diet (TestDiet 58V8, 35.8% CHO, 18.1% PRO, 46.1% FAT, semi-purified) from TestDiet (Richmond, IN) for eight weeks prior to the initiation of hormone treatment, and remained on HFD for another 25 days until the end of the experiment. OVX surgeries were conducted under 2% isoflurane and mice received Carprofen (0.074 mg/kg) and buprenorphine (0.1mg/kg) at the time of the surgery, with both drugs administered again at the same dose 24 hours after surgery. For hormone treatment, mice were implanted with osmotic mini pumps (as outlined below) that delivered vehicle (propylene glycol, n=6), 17βE2 (5 μg/day, n=7) or DHED (5 μg/day, n=7) treatment for 25 days. We conducted power analysis to determine the sample size required to detect an effect of 17βE2 treatment on body weight, which we considered the primary variable and a composite factor that reflects a change in energy balance with treatment. Previous studies haved reported strong effects of 17βE2 treatment on body weight changes after four weeks in the context of HFD treatment ^16,17,36^ and power analysis (Power at 0.8 and significance at α=0.05) using the mean body weight change and pooled standard deviation from previous studies^16^ indicated a small sample size (3-4 per group) was required to detect a similar magnitude change^16^. We allowed seven animals per group (six in controls) to account for any loss of animals over the experiment and to detect any weaker effects that may occur with other outcomes or DHED treatment. Mice were weighed weekly while on a HFD, and then every second day while receiving hormone treatment. Body weight and food intake were measured from four days post surgery so that any food or weight changes were not a consequence of acute adverse responses to surgery. Glucose tolerance tests (GTTs) were performed after 23 days of pump insertion and mice were euthanised via transcardial perfusion with 4% paraformaldehyde (PFA) for tissue collection after a total of 25 days of pump insertion.

**Fig. 1:**
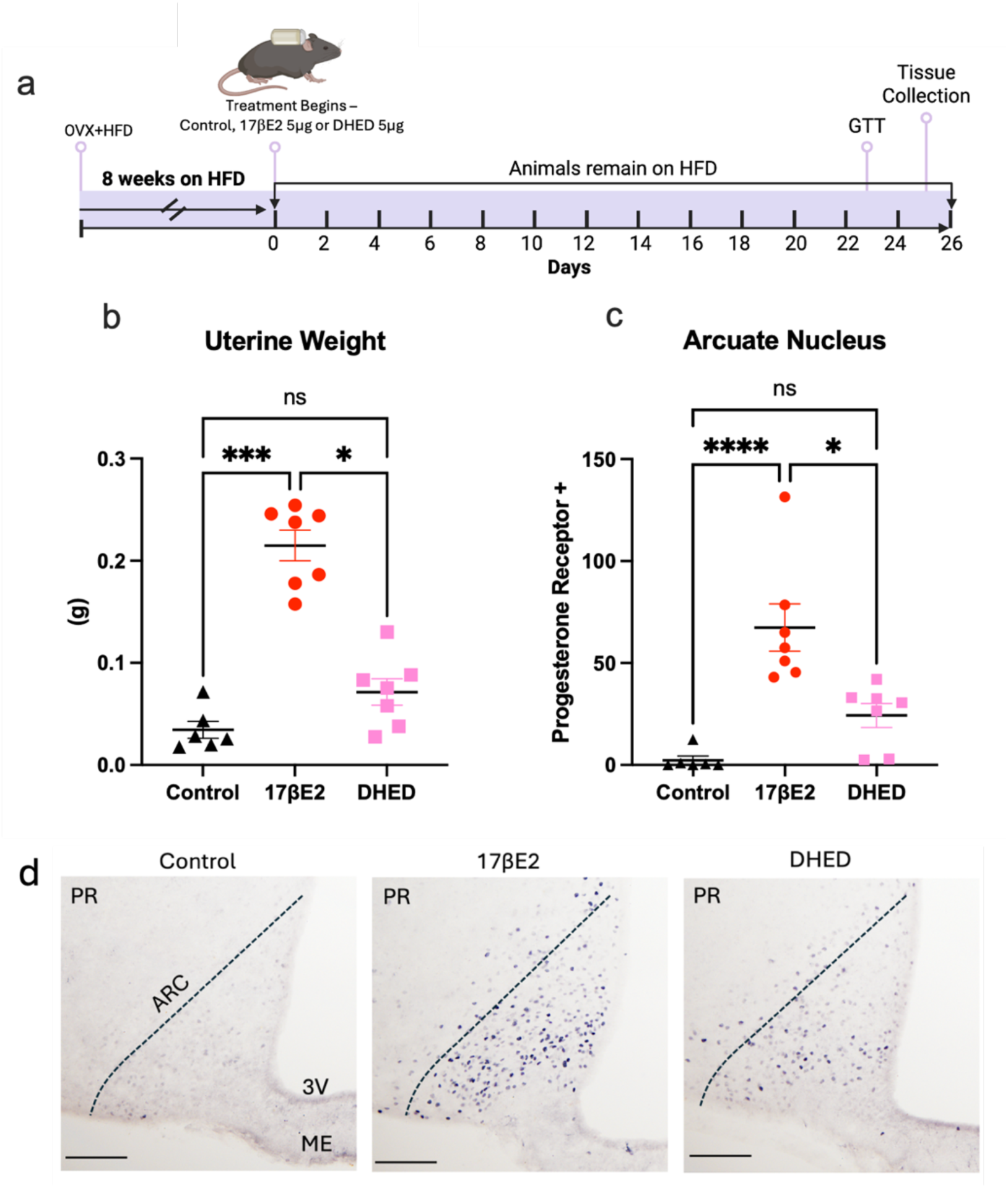
Experiment 1. DHED does not increase uterine weight and influences progesterone receptor expression in the ARC in some animals. **a**, timeline: Animals were ovariectomised and placed on a 45% HFD for eight weeks, then osmotic mini pumps delivered treatment for 25 days. n=6 for vehicle treatment, n =7 for 5 μg/day 17βE2 treatment, n=7 for 5 μg/day DHED treatment. b, Uterine weight for each animal c, Progesterone receptor (PR) abundance in the ARC d, Representative images of PR abundance: vehicle control, 17βE2 or DHED treatment. *p <0.05, ***p =0.001, ****p <0.0001, (Dunn’s post hoc test). Scale bar =100µm. Error bars show mean ± SEM. ARC= arcuate nucleus, 3V= third ventricle, ME=median eminence.

**Figure 2:**
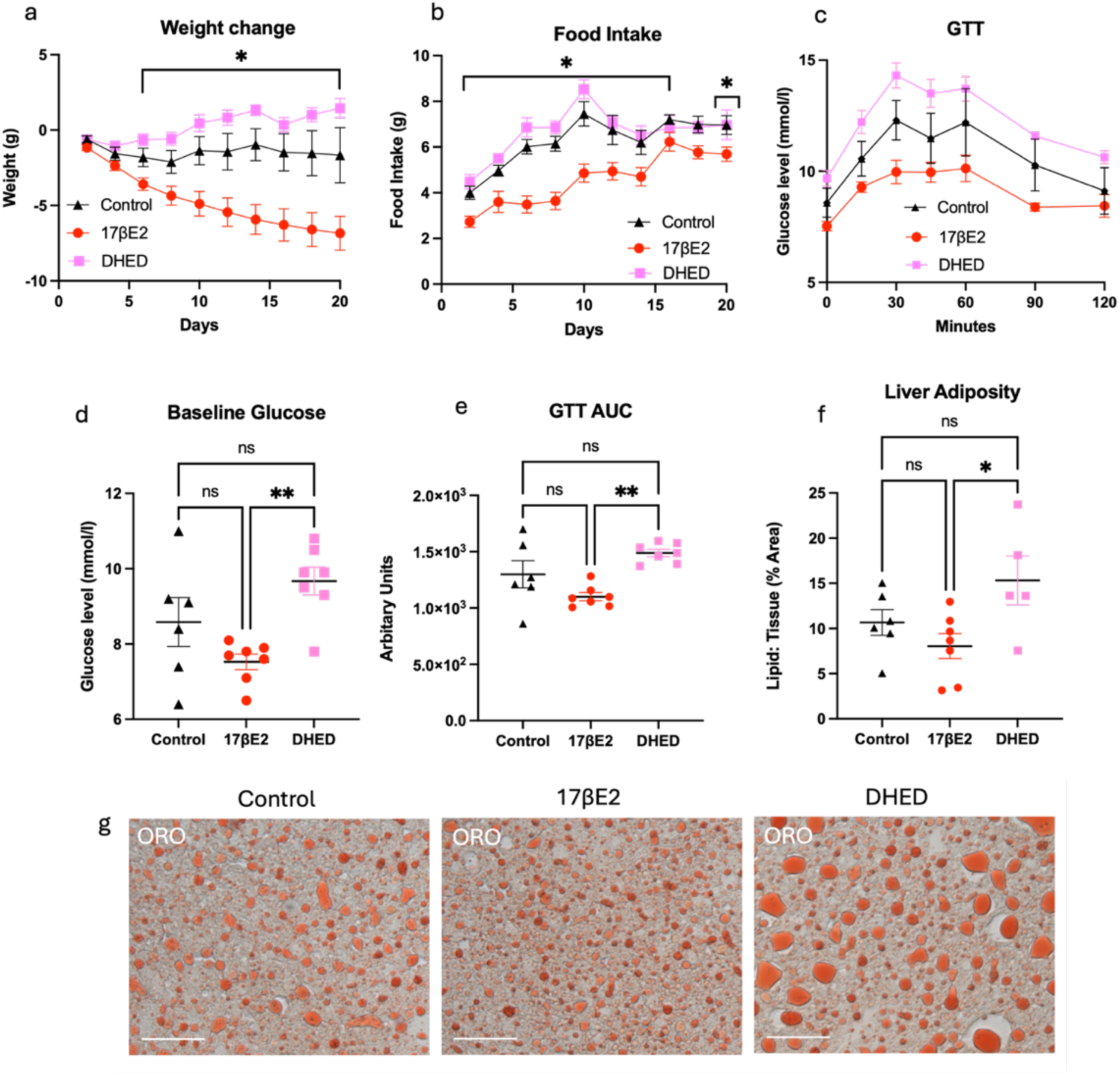
17βE2, but not DHED treatment, significantly improves metabolic parameters. **a,** Body weight change after initiation of hormone treatment (17βE2 or DHED) following eight weeks on HFD **b,** Food consumption, **c,** Glucose tolerance **d,** Baseline glucose levels, **e,** area under the curve **f,** Liver lipid levels **g,** Representative images showing liver lipid levels in response to treatment following Oil Red O (ORO) staining. **a,b,c**: *p <0.05 effect of 17βE2 relative to controls at denoted time points (Fisher’s LSD post hoc test). **d,e,f**: *p <0.05, **p <0.01 (Dunn’s post hoc test). n=6-7 per group. Error bars show mean ± SEM. Scale bar =75 µm.

#### Experiment 2: Acute DHED dose-response study to assess central actions

C57BL/6J mice maintained on a normal chow diet (Lab Diet, Mouse Diet 9F, (Protein: 23%; Fat 22%; Carbohydrates: 55%) underwent OVX at 7-9 weeks of age and were allowed to recover for one week before being split into groups that received either control (n=6, sesame oil), 17βE2 (5 μg/ day, n=4) or one of three increasing doses of DHED (5 μg /day (n=5), 10 μg/day (n=4), 20 μg/day (n=5)) (**Fig 3a**). The effect of estrogen treatment on PR expression is very strong, with all treated animals typically having high levels compared to OVX animals (i.e. **Fig 1c**, **Fig 3c**) ^37^. Power analysis indicated a minimum sample size for statistical analysis is required (i.e. n=3 per group) to detect an effect of 17βE2 treatment on PR expression; we therefore allowed 4-6 animals per group to account for any loss during the experiment or additional variability within DHED treated animals. DHED or 17βE2 was dissolved in 3% ethanol in sesame oil and administered once daily for two days via a subcutaneous injection **(Fig 3a)**. On day two, animals were sacrificed via transcardial perfusion with 4% PFA six hours after injection, a timeframe that is sufficient to observe increases in PR expression in the hypothalamus following 17βE2 treatment ^38^.

**Figure 3:**
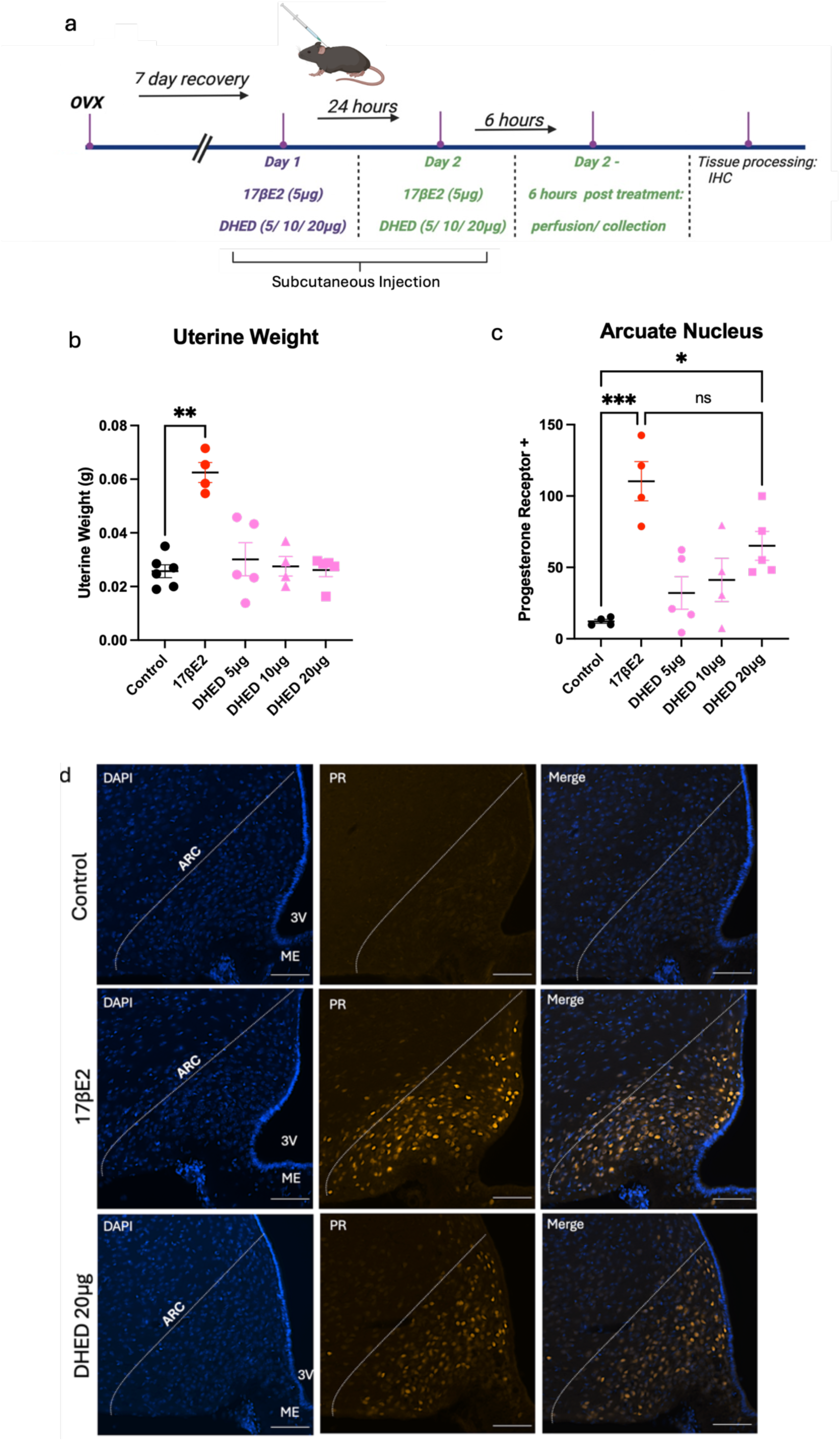
Experiment 2. **a,** timeline: Mice were ovariectomised, allowed to recover, then received either vehicle (n=6), 17βE2 (5 μg/day, n=4) or DHED (5 μg/day (n=5), 10 μg/day (n=4) or 20 μg/day (n=5)) once daily for two days via subcutaneous injection. Six hours post treatment on day two, animals were sacrificed and perfused with 4% PFA. **b,** Uterine weight after 48 hours of hormone treatment **c,** PR abundance in the ARC **d,** Representative images of PR immunoreactivity (orange) expression in the ARC (DAPI, PR only and merged channels) following control, 17βE2, or DHED 20 µg (see supplementary figure 1 for DHED 5 µg and 10 µg images). **p <0.01, ***p <0.001 effect of 17βE2 relative to controls, *p <0.05 effect of DHED 20 µg relative to controls (Dunn’s post hoc test). Error bars show mean ± SEM Scale bar = 100µm. ARC= arcuate nucleus, 3V= third ventricle, ME=median eminence.

#### Experiment 3: Prolonged DHED dose-response study to assess metabolic effects

C57BL/6J mice were ovariectomised (n=40) between 7-10 weeks of age (**Fig 4a**). Prior to OVX, mice were fed a normal chow diet (Teklad Global 18% Protein Rodent Diet #2918 (Protein: 18.6%; Fat 6.2%; Carbohydrate: 44.2%). Immediately following OVX, mice were placed on a 45% high fat diet (TestDiet 58V8, 35.8% CHO, 18.1% PRO, 46.1% FAT, semi-purified) from TestDiet (Richmond, IN) for four weeks prior to the initiation of hormone treatment and remained on this HFD for the remainder of the experiment. Mice then underwent surgery to insert osmotic mini pumps that delivered vehicle (polyethylene glycol, n=7), 17βE2 (5 μg/day, n=6) or one of three increasing doses of DHED (5 μg (n=6 per group), 10 μg (n=6 per group), 20 μg, n=7 per group) for 14 days. Mice were weighed weekly while on a HFD, and then every second day along with food intake while receiving hormone treatment. Body weight and food intake were measured for the first ten days. Glucose tolerance tests (GTTs) were performed on day 13 and mice were sacrificed via transcardial perfusion the following day.

**Fig 4:**
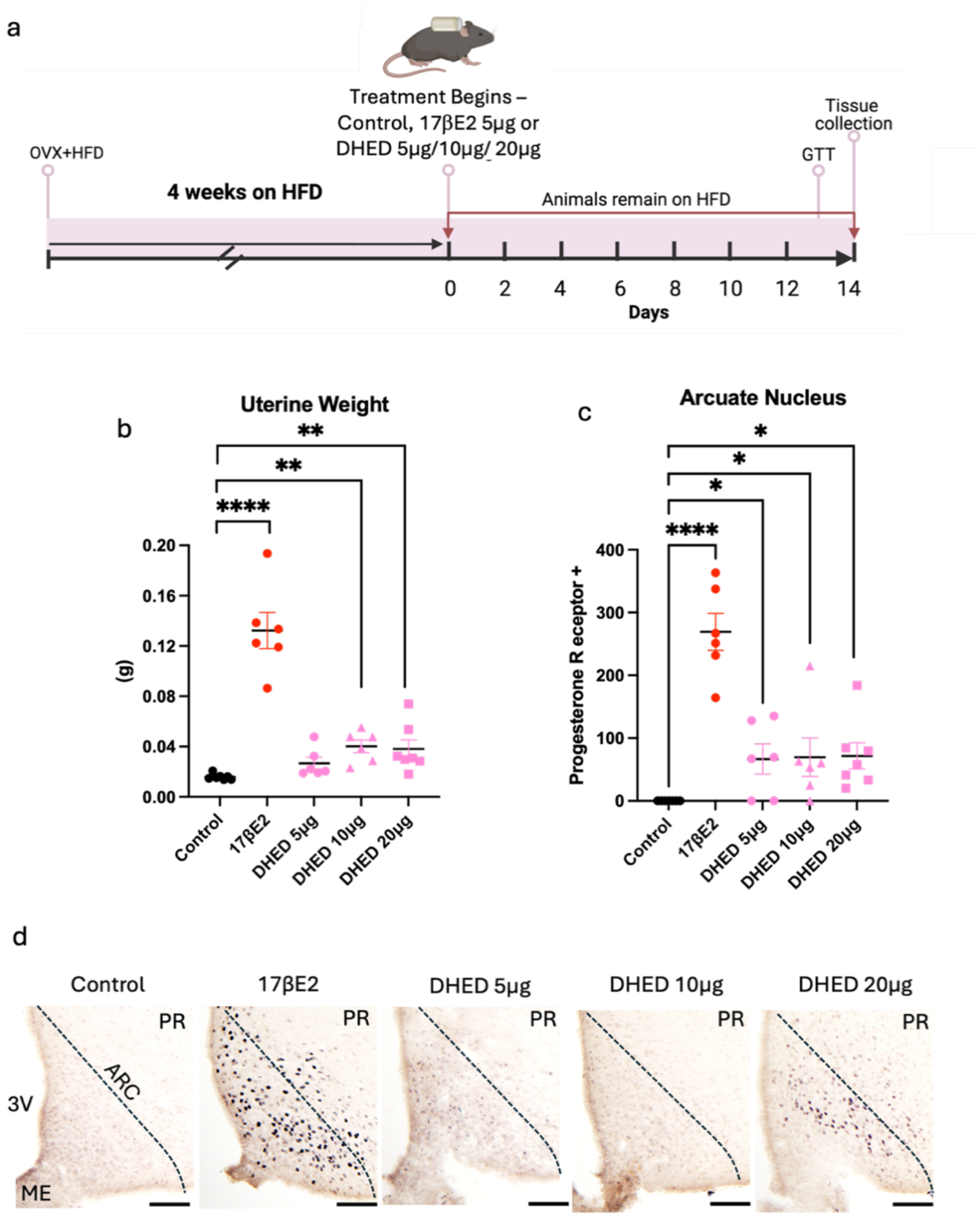
Experiment 3: DHED increases PR expression in the ARC. **a,** timeline: Animals were ovariectomised and placed on a 45% HFD for four weeks. Mice underwent a second surgery to insert osmotic mini pumps to deliver treatment (vehicle (n=7), 17βE2 (5 μg/day, n=6) or DHED (5μg/day, n=6, 10μg/day, n=6, 20 μg/day, n=7) for 14 days. At the end of the experiment all mice were perfused and tissues collected. **b,** Uterine weight in response to 17βE2 or increasing doses of DHED **c,** PR positive cells in the ARC. **d,** Representative images showing PR expression in the ARC following control, 17βE2 DHED 5 µg, DHED 10 µg, or DHED 20 µg. *p <0.05 effect of DHED 5, 10 or 20µg/day relative to controls, **p <0.01 effect of DHED 10 or 20µg/day relative to controls, ****p <0.0001 effect of 17βE2 relative to control. Error bars show mean ± SEM. Scale bar = 50µm. ARC= arcuate nucleus, 3V= third ventricle, ME=median eminence.

### Metabolic analyses

Mice were weighed weekly while on a HFD, and then every second day while osmotic mini pumps delivered treatments in experiments one and three. Food intake was assessed for pairs of mice in experiments one and three, with the average daily food intake value per cage used for analysis.

After mice had been on hormone treatment for 23 days (experiment one) or 13 days (experiment three), mice were fasted for 6 hours at 7-8am at the beginning of the light period (water access remained *ad libitum*) prior to intraperitoneal (ip) injection with 2g/kg D-glucose (n=6-7 per group). Blood glucose levels were measured with a glucose meter (Accu-CHEK, Roche) immediately pre-injection (time 0) and at 15-, 30-, 45-, 60-, 90-, and 120-minutes post-injection. For mice whose levels surpassed the Accu-CHEK meter range, the highest glucose reading possible of 33.3 mmol/l was used for that time point.

### Hormone administration via osmotic mini pumps

For experiments one and three, vehicle, 17βE2 and DHED treatments were delivered by subcutaneous infusion via osmotic mini pumps (Alzet, CA, USA, Model 2004, delivery rate 0.25 ± 0.05 μl/h for 25 days (experiment one), Alzet, CA, USA, Model 1002, delivery rate 0.25 ± 0.05 μl/h for 14 days (experiment three). In experiment one, 17βE2, DHED and vehicle solutions were dissolved and diluted in 100% propylene glycol so as to deliver 5 μg of either hormone per day . In experiment three, 17βE2, DHED and vehicle solutions were dissolved in 50% DMSO in polyethylene glycol 300 so as to dispense the required daily dosages. Pre-filled pumps were activated prior to insertion in a sterile saline solution (0.9%) in a 37°C water bath for 48 hours (experiment one) or overnight (experiment three). Mini pump insertion took place while animals were anesthetised under 2.5-3% isoflurane. Carprofen (0.074 mg/kg) and buprenorphine (0.1mg/kg) were administered at the time of the surgery, and both were administered again 24 hours after surgery.

### Euthanasia/tissue collection

At the end of each experiment mice were anesthetized with pentobarbital (150mg/kg) and upon loss of the pedal reflex, the uterus and a segment of liver were dissected. The animal was then perfused transcardially with 4% PFA in phosphate-buffered saline (PBS). Brains were dissected, post-fixed for 24 hours in 4% PFA in PBS, then immersed in 30% sucrose in PBS for another 24 hours. Brains were then frozen using dry ice and ethanol, embedded in optimal cutting temperature compound (OCT) and stored at -80°C.

### Experiments one and three: Immunohistochemistry for chromogenic labelling of PR

Brains were coronally sectioned in a series of four at thirty microns using a cryostat and stored in cryoprotectant at -20°C prior to processing. Two to three medial ARC sections were anatomically matched from each animal and were treated with 30% hydrogen peroxide solution in methanol and Tris Buffer Saline (TBS) prior to blocking (5% normal goat serum (NGS), TBS, 0.25% bovine serum albumin (BSA)). Sections were incubated at 4°C in a custom primary antibody against PR (1:2500, RC269, RRID: AB_2924988) for 60 hours^39,40^. Sections were then incubated for 60 minutes at room temperature (RT) in a secondary biotinylated immunoglobulin antibody (goat anti-rabbit, 1:250, Vector Laboratories Cat# BA-1000, RRID:AB_2313606) and then in Vectastain Elite Avidin-Biotin dissolved in antibody dilution solution (5ml/ml solution, Vector Laboratories) for 90 minutes at RT. Benzene derivative DAB (3,3′-Diaminobenzidine), nickel and glucose oxidase were used as a substrate to reveal a purple/brown derivative stain within the nucleus of positively stained cells for PR (n=6-7 per group). Sections were then washed, mounted, dried and dehydrated through a series of increasing concentrations of ethanol (50%, 70%, 90%, 100%) followed by 100% xylene and cover slipped using dibutylphthalate plasticizer xylene (DPX) mounting medium (Merck, #100579).

The number of PR positive cells in the ARC were quantified in two-three middle ARC sections (Paxinos coordinates: -1.46 to -1.70) from each mouse (n=6-7 per group). Sections were imaged on an Olympus 1501 microscope with a 10x objective. PR positive cell counts were automated with ImageJ software (experiment 1) or Qu Path (experiment 3) with constant thresholding throughout the analysis. The investigator was blinded to treatment group and conducted thresholding based on visible positive staining in the ARC. Briefly, the ImageJ protocol was as follows: The standardized ROI was outlined in each image (pixel size 0.2406 µm, 10x objective), the background was subtracted, and the particle analysis was conducted with the following measurements: rolling ball radius set to 250, pixel size 100-infinity, circularity 0.25-1. Only PR positive cells with a stained nucleus were included in counts. Qu Path analysis was performed with optical density sum; nucleus detection parameters were sigma 1.5 µm, background radius 8 µm, and area range 15–60 µm²; intensity thresholds were 0.2-0.25 with maximum background intensity 2 and split by shape enabled; cell expansion was set to 2 µm including the nucleus; and smooth boundaries with measurement recording were applied.

### Experiment two: Immunohistochemistry for fluorescent labelling of PR

Two to three middle ARC sections were anatomically matched from each animal and mounted onto super frost slides (n=4-5 per group). Slides were washed in PBS then blocked in a solution containing 10% NGS in 0.3% PBS-Triton for one hour, then incubated in a primary antibody against PR (1:1000, Abcam Cat# ab101688, RRID: AB_10715248) overnight in a dehumidified chamber. Sections were washed in PBS and then incubated for 2 hours at RT in an AlexaFluor647 conjugated secondary antibody (goat anti-rabbit, 1:1000, Invitrogen, Cat# A-11035, RRID: AB_2534093). Sections were washed once more in 1x PBS before being counterstained with nuclei marker (DAPI, 1:500) and cover-slipped using gold mounting media (Southern Biotech, 0100-01).

Images that encapsulated both sides of the ARC were collected using an Olympus 1501 microscope (20x), but unilateral (one side) counts were used for analysis. The average unilateral count from two to three middle ARC sections from each mouse were used. PR-positive cells were counted manually using the cell counter function in FIJI software. The investigator was blinded to treatment group. The ARC boundaries were defined consistently throughout the analysis. Only cells with PR-immunoreactive nucleus and a positive stain for the DAPI nuclear marker were included in counts.

### Liver Histology

Oil-red-O (ORO) staining was performed to assess lipid levels in liver samples as previously described ^41^. Twelve-micron liver sections were sectioned at -20°C on a cryostat and stored at -80°C (experiment one and three n=6-7 per group). ORO (2.5g) was dissolved in 400 ml of 99% isopropyl to make a stock solution, which was further diluted in distilled water to make a working solution. The working solution was filtered prior to use and sections were incubated in the working solution for 5 minutes, then rinsed in cold water for 30 minutes prior to sealing with nail varnish. Sections were imaged within 24 hours of staining on Olympus 1501 microscope (20x). The red lipid stain was quantified on ImageJ software using 5 regions of interest per animal and the investigator was blinded to treatment group. Staining is presented as a lipid to total tissue ratio. Thresholding was determined and remained consistent throughout the analysis.

### Statistical analyses

Statistics were performed using GraphPad Prism software (version 10). For each measured parameter, either an ordinary one-way ANOVA or Kruskal Wallis test was performed. A One-way ANOVA was used if the data were normally distributed and variance was similar between groups, but if these assumptions were not met, a Kruskal Wall test was applied. For body weight, food intake and glucose levels across the glucose tolerance test, a 2-way repeated measures ANOVA was performed, including body weight change from baseline, average daily food intake or glucose levels as the repeated dependent variable and treatment as an independent variable. Where a main group effect was observed, a Fisher’s LSD post-hoc test (or a Dunn’s post-hoc test for non-parametric comparisons) was performed to determine whether 17βE2 or DHED treatment groups differed to controls. Results are presented as mean ± SEM, unless otherwise stated.

## Results

### Experiment 1: DHED administration does not improve metabolic dysfunction in OVX mice but influences progesterone receptor expression in some animals

A previous study has shown that DHED treatment can provide benefits when provided at the same dose as 17βE2 in rodent models investigating depressive like behaviour and working memory, without influencing uterine weight ^42^. We therefore first tested DHED treatment at the same dose as 17βE2, with both hormones administered at 5 μg/day. After 25 days of treatment, 17βE2 treated animals had significantly increased uterine weights in comparison with vehicle treated animals, (**Fig 1b**, p <0.0001 overall effect of treatment, p <0.0001 for Fisher’s LSD post-hoc comparison between 17βE2 treated and control animals). DHED treated animals showed no significant increase in uterine weight compared to controls. DHED treated animals had much lighter uterine weights than 17βE2 treated animals (**Fig 1b,** p=0.013, comparison between 17βE2 and DHED treated animals).

PR within the ARC was significantly increased in response to 17βE2 treatment (**Fig 1c,d**, p <0.0001 overall effect of treatment, p <0.0001 for Fisher’s LSD post-hoc comparison between 17βE2 treated and control animals). While substantial PR abundance was detected in five of seven animals following DHED treatment, the observed increase did not reach significance compared to controls (**Fig 1c,d**, p =0.0735) and was significantly different to the observed increase with 17βE2 (**Fig 1c,d**, p =0.0210).

### 17βE2, but not DHED treatment, significantly reduces body weight and food intake in OVX mice exposed to a HFD

Almost immediately after the initiation of 17βE2 treatment, mice began to significantly reduce their body weight (**Fig 2a,** p =0.0003 overall effect of treatment across the measuring period, p <0.05 effect of 17βE2 in comparison to control animals at each time point after 4 days). DHED administration did not recapitulate this protective effect of reduced body weight, with DHED-treated individuals showing no significant difference in body weight compared to controls. Food intake was also significantly reduced almost immediately after the initiation of treatment in the 17βE2 group (**Fig 2b**, p <0.0001 overall effect of treatment, p <0.05 effect of 17βE2 relative to controls at denoted time points), but DHED did not influence food intake compared to controls.

### Glucose tolerance is not improved in response to either DHED or 17βE2 treatment

In contrast to the hormone’s effect on body weight and food intake, 17βE2 treatment did not significantly lower glucose levels compared to control animals (**Fig 2c,** p =0.0176 overall effect of treatment). There was a trend across the experiment for significantly improved glucose tolerance in 17βE2 treated mice, most notably thirty minutes after after glucose injection (p = 0.0535), however no significant difference was observed between groups at any timepoint including at the baseline (**Fig 2c,d**). While the glucose tolerance test, baseline glucose levels and area under the curve (AUC) showed an overall effect of treatment (**Fig 2c,d,e,** p <0.05), the post hoc analysis revealed this was likely driven by differences between hormone treated animals (p <0.01, effect of 17βE2 in comparison with DHED treated animals), as there were no differences between either 17βE2 or DHED treated animals in comparison with controls for any parameter. Taken together, these results illustrate that neither 17βE2 or DHED improves glucose control in these mice.

### Liver adiposity is not reduced in response to either 17βE2 or DHED treatment

17βE2 has previously been reported to reduce liver adiposity in OVX female rodents ^43^, however we did not observe a reduction in hepatic lipid levels in 17βE2 treated mice (**Fig 2f,g** p =0.2550 effect of 17βE2 in comparison to control animals). While we observed an overall effect of treatment (**Fig 2f,g**, p =0.0492), this is not explained by any comparison to control animals and rather is driven by differences between hormone-treated groups (p =0.0492 overall effect of treatment, p <0.05 comparison between 17βE2 and DHED animals). We also observed no change in lipid abundance in DHED-treated mice compared to controls (**Fig 2f,g).**

### Experiment 2: DHED increases progesterone receptor expression in the ARC in a dose dependant manner when delivered over 48 hours

We subsequently performed an acute study over 48 hours to ascertain the dose of DHED required to elicit similar hypothalamic responses as 17βE2. After two days of once daily delivery, none of the DHED doses influenced uterine weight compared to control animals (**Fig 3b**, overall effect of treatment p <0.05, effect of DHED treatment at any dose compared to controls p >0.05). In contrast, two days of 17βE2 treatment resulted in significantly heavier uterine weights than control animals (**Fig 3b**, p <0.01 effect of 17βE2).

### Acute administration of DHED over 48 hours increases the expression of PR in a dose dependent manner

When administered acutely at the same dose of 5 µg as used in the previous experiment, 17βE2 significantly increased PR expression after 48 hours (**Fig 3c**, p <0.01 overall treatment effect, ***p < 0.001 effect of 17βE2). However, 48 hours of DHED treatment at the same dosage as 17βE2 or administered at a two-fold higher dose of 10 µg/day failed to significantly increase PR expression in this acute treatment setting (p >0.05). A significant increase in the number of cells expressing PR was only observed when DHED was administered at a four-fold increase of 20 µg/day (**Fig 3c,d**, p =0.026 effect of 20 µg/day DHED treatment relative to controls). There was also no statistical difference observed in number of PR+ cells in the ARC between 17βE2 and DHED administered at 20 µg/day.

### Experiment 3: DHED increases PR expression in the ARC but does not influence metabolic health at any dose

Given that a higher dose of acutely administered DHED (20 µg) was required to see estradiol-like actions in the brain, we carried out a longer-term study with a range of increasing DHED doses. After 14 days of 17βE2 treatment, animals had significantly increased uterine weights in comparison with vehicle-treated animals (**Fig 4b**, p <0.0001, overall effect of treatment). DHED at 5µg/day had no significant effect on uterine weight compared to controls but showed a pattern that was reflective of that seen in the first experiment (**Fig 4b**, p =0.0963). Animals treated with a higher dose of DHED at either 10 or 20µg/day had increased uterine weights relative to controls (p =0.0042, p =0.0099 effect of 10 or 20µg/day respectively relative to controls), but the 20µg/day group were significantly lighter than 17βE2 treated animals (p <0.05).

PR expression was robustly increased in response to chronic 17βE2 treatment (**Fig 4c,** p <0.001 overall effect of treatment, p <0.0001 effect of 17βE2 relative to control animals). Chronic DHED treatment also increased PR expression at all of the three doses tested (**Fig 4c,d**, p <0.05 effect of each dose in comparison to control animals), although to a lower degree.

### Increasing DHED dosages does not improve body weight or food intake

Over the first week of 17βE2 treatment, mice began to reduce their body weight (**Fig 5a),** which was consistent with our previous experiment (p <0.001 overall effect of treatment across the measuring period). DHED treatment did not recapitulate this protective effect of reduced body weight at any dose, with DHED-treated individuals showing no significant difference in body weight compared to controls. Food intake was also significantly reduced almost immediately after the initiation of treatment in the 17βE2 group (**Fig 5b,** p <0.01 overall effect of treatment across the measuring period) but none of the doses of DHED (5, 10, 20 µg) significantly influenced food intake in this cohort.

**Fig 5:**
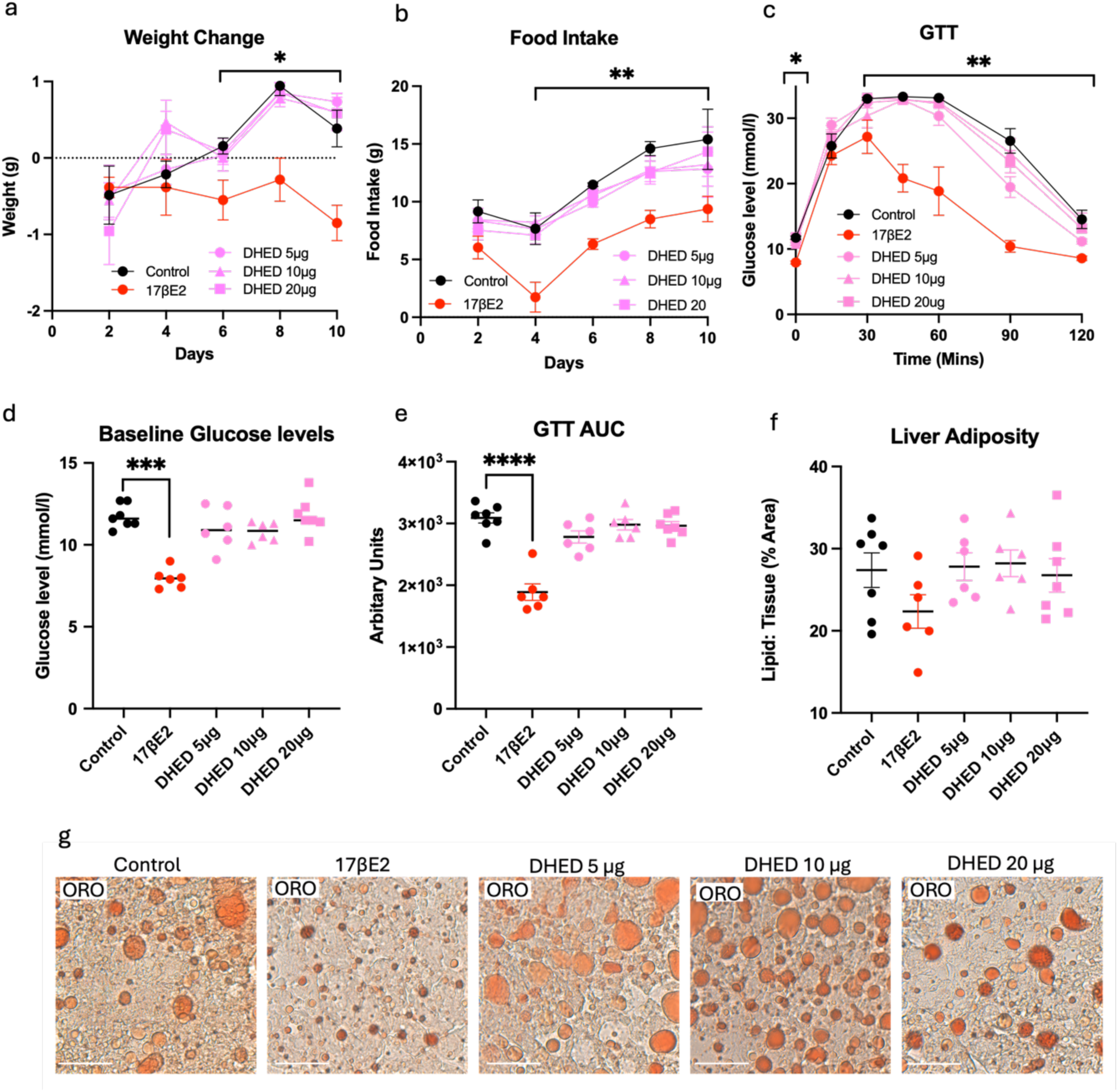
Increasing doses of DHED does not improve metabolic health in OVX obese mice. **a,** Body weight change in response to 17βE2 or DHED treatment. **b,** Food intake **c,** Glucose tolerance **d,** Baseline glucose levels **e,** area under the curve. **f,** Liver lipid levels **g,** Representative images showing liver lipid levels following ORO staining. **a,b,c,** *p <0.05, **p <0.01, effect of 17βE2 relative to controls from Fisher’s LSD post hoc test at each timepoint. **d,e** ***p <0.001, ****p <0.0001 effect of 17βE2 relative to controls determined from Dunn’s post hoc test. Error bars show mean ± SEM. n=6-7 per group. Scale bar =75 µm.

### DHED treatment does not improve glucose tolerance or liver adiposity at increasing doses

17βE2 significantly improved glucose tolerance compared to controls, with animals in this group showing a reduced glucose concentration at every time point from 30 minutes post glucose injection (**Fig 5c**, p <0.0001 overall effect of treatment), a reduction in baseline glucose levels (**Fig 5d,** p =0.0011 overall effect of treatment), and a significantly smaller AUC (**Fig 5e**, p <0.01 overall effect of treatment). In contrast, DHED treatment at either 5, 10 or 20 µg failed to have any effect on fasting glucose levels, glucose tolerance or AUC (**Fig 5c-e**). Liver adiposity was also not significantly reduced in response to either 17βE2 or any dose of DHED treatment, as similarly to the first experiment, no changes in hepatic lipid levels or lipid abundance were measured in any treatment group (**Fig 5f,g**, overall effect of treatment p= 0.2780).

## Discussion

This study sought to determine whether administration of DHED, a brain-targeting prodrug of 17βE2, could recapitulate the protective metabolic effects of 17βE2 treatment in OVX mice perturbed by a HFD. Confirming previous reports, we found that 17βE2 significantly improved metabolic parameters in obese OVX female mice receiving HFD treatment ^16,17,36,44^. In contrast, despite demonstrating actions in the brain, DHED administration failed to improve a single metabolic parameter across either of the metabolic experiments at any dose tested, similar to a recent report in aged mice on a standard rodent chow diet ^45^. Taken together, our results suggest that exclusive targeting of the brain with 17βE2 via DHED treatment is insufficient to improve metabolic health in OVX mice.

The dose of 17βE2 used was equivalent to that used in previous studies that have shown beneficial effects of 17βE2 on metabolic health ^16,17,36,46^. Previous studies investigating DHED’s capacity to provide benefits in cognitive decline, hot flush and depression have suggested that the same dose of the prodrug as 17βE2 is sufficient to alleviate symptoms ^31,32,34^. Across our study, direct administration of 17βE2 significantly increased uterine weight compared to controls as this increases the presence of this hormone in the circulation. In contrast, short-term or low-dose chronic DHED administration had no effect on uterine weight, despite showing evidence for actions in the brain. This suggests that DHED actions were restricted to the brain in this context. However, higher doses of DHED delivered over 14 days resulted in an approximately two-fold increase in uterine weight relative to controls, indicating the estrogenic actions of DHED are not restricted to the brain at higher doses. However, the increase was modest when compared with the proliferative effects of 17βE2, with animals in this treatment group showing an approximately 6-fold increase in uterine weight. Similarly, post hoc statistical analyses revealed a significant difference between 17βE2 (5µg) and the 20µg DHED group in our third experiment. Given that no previous study has reported an increase on uterine weight and previous extensive characterization has failed to detect 17βE2 in peripheral tissues ^32,45,47^, it is possible that perturbation by a HFD may partially explain this increase in uterine weight. HFD induces chronic inflammation and an increase in oxidative stress in the brain, including within the ARC ^48,49^. Resulting neuroinflammation can lead to a wide array of pathophysiological effects in the brain, including mitochondrial dysfunction and neuronal death ^50^. Given that neuroinflammatory populations can also alter blood brain barrier permeability, it is plausible that a HFD is altering typical brain signalling actions and is causing some leakage of DHED from the brain back into circulation, particularly at high concentrations ^51^. Unfortunately it was not possible to measure 17βE2 in this study, with more commercially available ELISAs lacking sufficient specificity. Future studies with direct comparisons between normal chow and HFD will help to elucidate this finding.

In all three experiments, we used PR expression as a reporter and positive control for estrogen receptor actions within the brain ^52–54^. Previously, the only hypothalamic (or brain) nucleus that has been probed for PR expression following DHED treatment is the preoptic area (POA), which was conducted in OVX rats ^32,47^. Here, we show that DHED treatment only increases PR expression within the ARC when delivered at a four-fold increased dose of 17βE2 in OVX mice when delivered acutely. In our first study over 25 days, we show that a similar dose of DHED does not increase PR to the same levels as 17βE2, whereas when delivered for 14 days, we show a ceiling effect with increasing doses of the prodrug, as a similar, two-fold increased or four fold increased dose of DHED increased PR expression to a similar degree, but failed to reach levels observed with 17βE2 treatment. While it is likely that brain-specific delivery of 17βE2 is not sufficient to improve metabolic health independant of peripheral estrogen actions, further extensive anaylsis of how DHED is converted into 17βE2 within the medio-basal located ARC (and throughout the hypothalamus) is required before this conclusion can be determined.

To date, the sole other study investigating DHED in the context of metabolic dysfunction showed that body weight or glucose tolerance are not improved in normal chow mice, even when DHED is administered at a significantly higher dose than in our study ^45^. In that study, 11.5-month old OVX mice were fed a standard chow diet and treated with DHED for 4 months and these animals showed no differences in body weight compared to controls. Surprisingly, DHED-treated animals also had impaired glucose tolerance when applied at high doses. However, these tests were conducted in aged mice after an extended period of behavioural testing, and the use of standard chow meant that females were not suffering impaired metabolic dysfunction from a HFD, which is the context where metabolic effects of 17βE2 have typically been explored. Moreover, since in that study no 17βE2 treatment group was included, it is not possible to make comparisons of the effectiveness of DHED to 17βE2 in modulating metabolic health in either the presence or absence of obesity.

The brain is a target of 17βE2-mediated metabolic actions and can modulate energy homeostasis by reducing food intake through actions on specific neuronal populations within the hypothalamus ^55,56^. 17βE2 can influence the secretion of anorexigenic neuropeptides which in turn reduce food intake and promote satiety, including pomc, leptin and insulin, all of which have receptors within the ARC of the hypothalamus ^57–59^. Ablation of ERs throughout the CNS, both from within specific cellular populations (including *POMC)* and regions (VMHvl, ARC) initiates many metabolic disturbances including hyperphagia, reduced energy expenditure, weight gain and reduced activity ^25,26,29^. This would suggest that 17βE2’s actions within the brain are necessary to drive protective effects on metabolic health. However, our findings presented here (in addition to the other report investigating DHED’s effects on body weight and glucose tolerance) suggest that central estradiol signalling, while necessary to promote protective metabolic health, appears insufficient to achieve this protection when present independent of peripheral estrogen actions ^45^.

Our results could point to an important role of peripheral estrogen receptors in metabolically active tissues, which therefore cannot be overlooked in their role in regulating metabolic function as inidicated by previous tissue-specific knockout studies. Ablation of peripheral ERs disrupts metabolic health, by altering triglyceride levels (liver-specific knockout models), enhancing adipocyte mass (adipocyte-specific knockouts) and disrupting bone turnover (global knockouts) ^60–63^. It is therefore likely that exclusive targeting of the brain with 17βE2 is insufficient to independently drive protective effects on metabolism, because these peripheral actions are required concurrently with brain actions. This may explain why the metabolic improvements we observed with 17βE2 were absent in DHED treated animals, as the peripheral actions of 17βE2 may be required to cause these protective effects.

To conclude, our investigations reveal that exclusive targeting of the brain with 17βE2 via the DHED prodrug fails to offer any benefit on metabolic health in obese, OVX mice, despite evidence that this treatment is inducing some aspects of estrogen signalling within the ARC of the hypothalamus. Future directions focusing on alternatives to traditional MHT should include integrated approaches between 17βE2’s actions in the brain and periphery, as our results suggest that peripheral estrogen signalling is required to initiate 17βE2-mediated protective effects on metabolic health.

## Grants/Fellowships

The authors would like to express their gratitude to the Elman Poole Foundation, the British Society for Neuroendocrinology, the Hope Foundation for Research on Ageing and the University of Otago’s Doctoral Scholarship for their support to CC.

## Acknowledgements

The authors would like to thank members of the Garratt laboratory who assisted with glucose tolerance tests and the animal technicians in the Biomedical Research Facilities at both the University of Otago and University of California, Los Angeles.

## Disclosures

KP-T is an inventor in the DHED patents and had equity interest in Agypharma LLC that licensed the approach for commercial development.

## Appendix

**Supplementary Figure 1:**
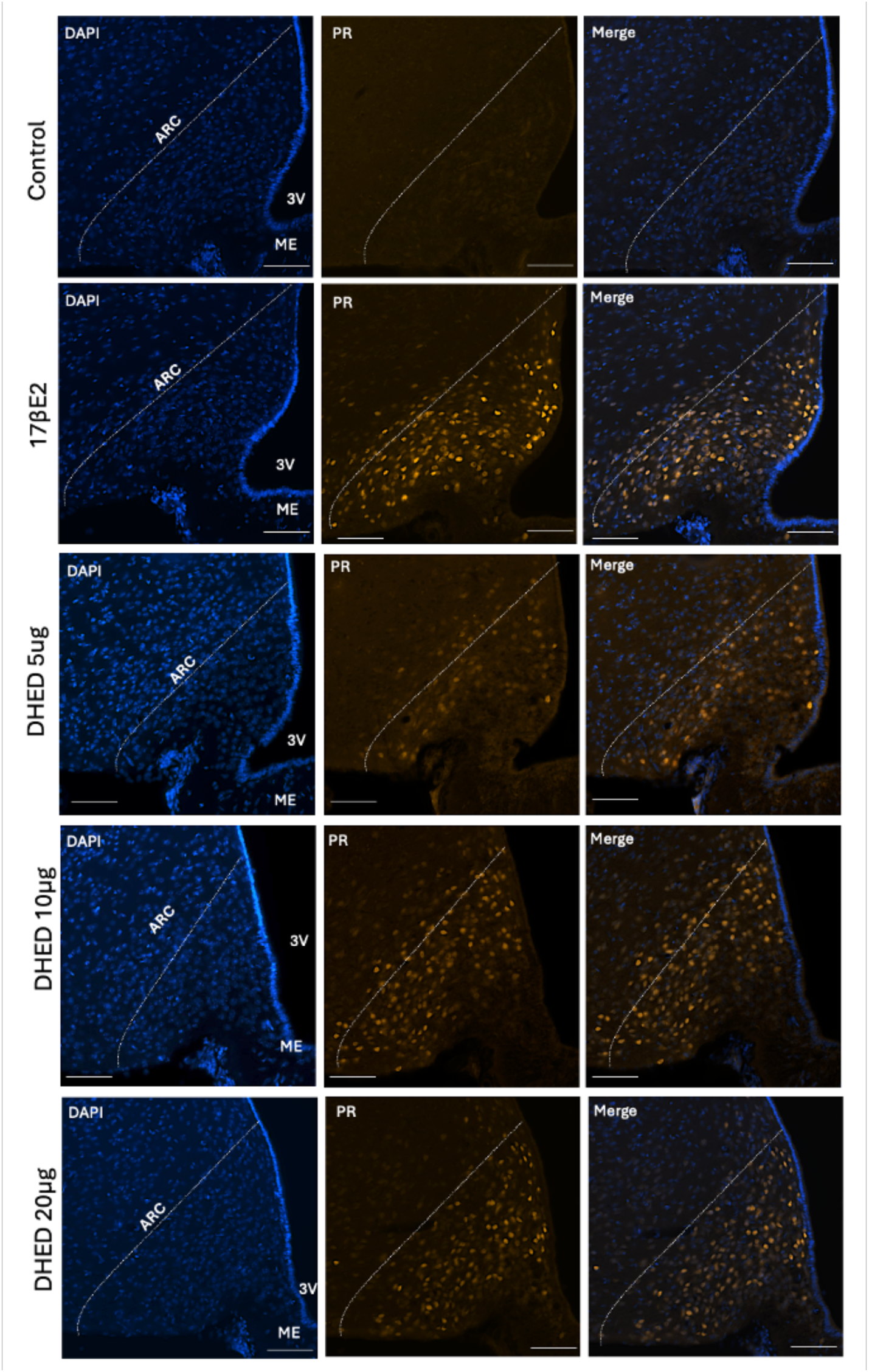
Experiment 2: **a,** Representative images of PR immunoreactivity (orange) expression in the ARC (DAPI, PR only and merged channels) following control, 17βE2, DHED 5 µg, DHED 10 µg or DHED 20 µg. Overall effect of treatment p <0.05 for **b** and **c** (Kruskal-Wallis test), **p <0.01, ***p <0.001 effect of 17βE2 relative to controls, *p <0.05 effect of DHED 20 µg relative to controls (Dunn’s post hoc test). Error bars show mean ± SEM Scale bar = 50µm.

## References

1. Abdelmonem AH. Is There any Mean to Postpone The Menopausal Ovarian Senescence? Int J Fertil Steril. 2020;13(4):346–347. doi:10.22074/ijfs.2020.5797

2. Burger HG, Hale GE, Dennerstein L, Robertson DM. Cycle and hormone changes during perimenopause: the key role of ovarian function. Menopause. 2008;15(4):603. doi:10.1097/gme.0b013e318174ea4d

3. Polo-Kantola P. Dealing with Menopausal Sleep Disturbances. Sleep Med Clin. 2008;3(1):121–131. doi:10.1016/j.jsmc.2007.10.006

4. Bruce D, Rymer J. Symptoms of the menopause. Best Pract Res Clin Obstet Gynaecol. 2009;23(1):25–32. doi:10.1016/j.bpobgyn.2008.10.002

5. Kodoth V, Scaccia S, Aggarwal B. Adverse Changes in Body Composition During the Menopausal Transition and Relation to Cardiovascular Risk: A Contemporary Review. Womens Health Rep. 2022;3(1):573–581. doi:10.1089/whr.2021.0119

6. Akahoshi M, Soda M, Nakashima E, Shimaoka K, Seto S, Yano K. Effects of Menopause on Trends of Serum Cholesterol, Blood Pressure, and Body Mass Index. Circulation. 1996;94(1):61–66. doi:10.1161/01.CIR.94.1.61

7. Vishvanath L, Gupta RK. Contribution of adipogenesis to healthy adipose tissue expansion in obesity. J Clin Invest. 2019;129(10):4022–4031. doi:10.1172/JCI129191

8. Mattsson LA, Skouby S, Rees M, et al. Efficacy and tolerability of continuous combined hormone replacement therapy in early postmenopausal women. Menopause Int. 2007;13(3):124–131. doi:10.1258/175404507781605596

9. Marsh ML, Oliveira MN, Vieira-Potter VJ. Adipocyte Metabolism and Health after the Menopause: The Role of Exercise. Nutrients. 2023;15(2):444. doi:10.3390/nu15020444

10. Camon C, Garratt M, Correa SM. Exploring the effects of estrogen deficiency and aging on organismal homeostasis during menopause. Nat Aging. 2024;4(12):1731–1744. doi:10.1038/s43587-024-00767-0

11. Chen Y, Heiman ML. Increased weight gain after ovariectomy is not a consequence of leptin resistance. Am J Physiol-Endocrinol Metab. 2001;280(2):E315–E322. doi:10.1152/ajpendo.2001.280.2.E315

12. Rogers NH, Perfield JW II, Strissel KJ, Obin MS, Greenberg AS. Reduced Energy Expenditure and Increased Inflammation Are Early Events in the Development of Ovariectomy-Induced Obesity. Endocrinology. 2009;150(5):2161–2168. doi:10.1210/en.2008-1405

13. MacLean PS, Giles ED, Johnson GC, et al. A Surprising Link Between the Energetics of Ovariectomy-induced Weight Gain and Mammary Tumor Progression in Obese Rats. Obesity. 2010;18(4):696–703. doi:10.1038/oby.2009.307

14. Wang Y, Wang Y, Liu L, Cui H. Ovariectomy induces abdominal fat accumulation by improving gonadotropin-releasing hormone secretion in mouse. Biochem Biophys Res Commun. 2022;588:111–117. doi:10.1016/j.bbrc.2021.12.039

15. Garratt M, Lagisz M, Staerk J, et al. Sterilization and contraception increase lifespan across vertebrates. Nature. Published online December 10, 2025:1–9. doi:10.1038/s41586-025-09836-9

16. Stubbins RE, Holcomb VB, Hong J, Núñez NP. Estrogen modulates abdominal adiposity and protects female mice from obesity and impaired glucose tolerance. Eur J Nutr. 2012;51(7):861–870. doi:10.1007/s00394-011-0266-4

17. Fuller KNZ, McCoin CS, Von Schulze AT, Houchen CJ, Choi MA, Thyfault JP. Estradiol treatment or modest exercise improves hepatic health and mitochondrial outcomes in female mice following ovariectomy. Am J Physiol - Endocrinol Metab. 2021;320(6):E1020–E1031. doi:10.1152/ajpendo.00013.2021

18. Magnusson C, Baron JA, Correia N, Bergström R, Adami HO, Persson I. Breast-cancer risk following long-term oestrogen- and oestrogen-progestin-replacement therapy. Int J Cancer. 1999;81(3):339–344. doi:10.1002/(sici)1097-0215(19990505)81:3%3C339::aid-ijc5%3E3.0.co;2-6

19. Rodriguez AC, Blanchard Z, Maurer KA, Gertz J. Estrogen Signaling in Endometrial Cancer: a Key Oncogenic Pathway with Several Open Questions. Horm Cancer. 2019;10(2-3):51–63. doi:10.1007/s12672-019-0358-9

20. Liu T, Xu Y, Yi CX, Tong Q, Cai D. The hypothalamus for whole-body physiology: from metabolism to aging. Protein Cell. 2022;13(6):394–421. doi:10.1007/s13238-021-00834-x

21. Xu Y, Nedungadi TP, Zhu L, et al. Distinct hypothalamic neurons mediate estrogenic effects on energy homeostasis and reproduction. Cell Metab. 2011;14(4):453–465. doi:10.1016/j.cmet.2011.08.009

22. Mauvais-Jarvis F, Clegg DJ, Hevener AL. The Role of Estrogens in Control of Energy Balance and Glucose Homeostasis. Endocr Rev. 2013;34(3):309–338. doi:10.1210/er.2012-1055

23. Simerly RB, Young BJ. Regulation of estrogen receptor messenger ribonucleic acid in rat hypothalamus by sex steroid hormones. Mol Endocrinol Baltim Md. 1991;5(3):424–432. doi:10.1210/mend-5-3-424

24. Yokosuka M, Okamura H, Hayashi S. Postnatal development and sex difference in neurons containing estrogen receptor-α immunoreactivity in the preoptic brain, the diencephalon, and the amygdala in the rat. J Comp Neurol. 1997;389(1):81–93. doi:10.1002/(SICI)1096-9861(19971208)389:1%3C81::AID-CNE6%3E3.0.CO;2-A

25. Camon C, Prescott M, Neyt C, et al. Systemic metabolic benefits of 17α-estradiol are not exclusively mediated by ERα in glutamatergic or GABAergic neurons. GeroScience. Published online May 22, 2024. doi:10.1007/s11357-024-01192-2

26. Xu Y, Nedungadi TP, Zhu L, et al. Distinct Hypothalamic Neurons Mediate Estrogenic Effects on Energy Homeostasis and Reproduction. Cell Metab. 2011;14(4):453–465. doi:10.1016/j.cmet.2011.08.009

27. Jais A, Brüning JC. Arcuate Nucleus-Dependent Regulation of Metabolism—Pathways to Obesity and Diabetes Mellitus. Endocr Rev. 2021;43(2):314–328. doi:10.1210/endrev/bnab025

28. He Y, Xu P, Wang C, et al. Estrogen receptor-α expressing neurons in the ventrolateral VMH regulate glucose balance. Nat Commun. 2020;11(1):2165. doi:10.1038/s41467-020-15982-7

29. van Veen JE, Kammel LG, Bunda PC, et al. Hypothalamic estrogen receptor alpha establishes a sexually dimorphic regulatory node of energy expenditure. Nat Metab. 2020;2(4):351–363. doi:10.1038/s42255-020-0189-6

30. Beall C, Ashford ML, McCrimmon RJ. The physiology and pathophysiology of the neural control of the counterregulatory response. Am J Physiol-Regul Integr Comp Physiol. 2012;302(2):R215–R223. doi:10.1152/ajpregu.00531.2011

31. Prokai L, Nguyen V, Szarka S, et al. The prodrug DHED selectively delivers 17β- estradiol to the brain for treating estrogen-responsive disorders. Sci Transl Med. 2015;7(297):297ra113. doi:10.1126/scitranslmed.aab1290

32. Merchenthaler I, Lane M, Sabnis G, et al. Treatment with an orally bioavailable prodrug of 17β-estradiol alleviates hot flushes without hormonal effects in the periphery. Sci Rep. 2016;6(1):30721. doi:10.1038/srep30721

33. Tschiffely AE, Schuh RA, Prokai-Tatrai K, Prokai L, Ottinger MA. A comparative evaluation of treatments with 17β-estradiol and its brain-selective prodrug in a double-transgenic mouse model of Alzheimer’s disease. Horm Behav. 2016;83:39–44. doi:10.1016/j.yhbeh.2016.05.009

34. Prokai-Tatrai K, Prokai L. A Novel Prodrug Approach for Central Nervous System-Selective Estrogen Therapy. Molecules. 2019;24(22):4197. doi:10.3390/molecules24224197

35. Quadros PS, Wagner CK. Regulation of Progesterone Receptor Expression by Estradiol Is Dependent on Age, Sex and Region in the Rat Brain. Endocrinology. 2008;149(6):3054–3061. doi:10.1210/en.2007-1133

36. Camporez JPG, Jornayvaz FR, Lee HY, et al. Cellular Mechanism by Which Estradiol Protects Female Ovariectomized Mice From High-Fat Diet-Induced Hepatic and Muscle Insulin Resistance. Endocrinology. 2013;154(3):1021–1028. doi:10.1210/en.2012-1989

37. Brown T, Clark A, MacLusky N. Regional sex differences in progestin receptor induction in the rat hypothalamus: effects of various doses of estradiol benzoate. J Neurosci. 1987;7(8):2529–2536.

38. Parsons B, Rainbow TC, Pfaff DW, McEwen BS. Oestradiol, sexual receptivity and cytosol progestin receptors in rat hypothalamus. Nature. 1981;292(5818):58–59. doi:10.1038/292058a0

39. Ruddenklau A, Glendining K, Prescott M, Campbell RE. Validation of a new Custom Polyclonal Progesterone Receptor Antibody for Immunohistochemistry in the Female Mouse Brain. J Endocr Soc. 2023;7(10):bvad113. doi:10.1210/jendso/bvad113

40. Lott EEA, Prescott M, Potapov K, Handelsman DJ, Glendining KA, Campbell RE. Forebrain AR Deletion Restores PR Expression but not Reproduction in Prenatally Androgenized Female Mice. Endocrinology. 2025;166(12):bqaf161. doi:10.1210/endocr/bqaf161

41. Mehlem A, Hagberg CE, Muhl L, Eriksson U, Falkevall A. Imaging of neutral lipids by oil red O for analyzing the metabolic status in health and disease. Nat Protoc. 2013;8(6):1149–1154. doi:10.1038/nprot.2013.055

42. Prokai-Tatrai K, Prokai L. A Novel Prodrug Approach for Central Nervous System-Selective Estrogen Therapy. Molecules. 2019;24(22):4197. doi:10.3390/molecules24224197

43. Tian Y, Xie Y, Hong X, Guo Z, Yu Q. 17β-Estradiol protects female rats from bilateral oophorectomy-induced nonalcoholic fatty liver disease induced by improving linoleic acid metabolism alteration and gut microbiota disturbance. Heliyon. 2024;10(7):e29013. doi:10.1016/j.heliyon.2024.e29013

44. Handgraaf S, Dusaulcy R, Visentin F, Philippe J, Gosmain Y. 17-β Estradiol regulates proglucagon-derived peptide secretion in mouse and human α- and L cells. JCI Insight. 3(7):e98569. doi:10.1172/jci.insight.98569

45. Salinero AE, Abi-Ghanem C, Venkataganesh H, et al. Treatment with brain specific estrogen prodrug ameliorates cognitive effects of surgical menopause in mice. Horm Behav. 2024;164:105594. doi:10.1016/j.yhbeh.2024.105594

46. Handgraaf S, Dusaulcy R, Visentin F, Philippe J, Gosmain Y. 17-**β** Estradiol regulates proglucagon-derived peptide secretion in mouse and human **α**- and L cells. JCI Insight. 2018;3(7). doi:10.1172/jci.insight.98569

47. Prokai L, Nguyen V, Szarka S, et al. The prodrug DHED selectively delivers 17β-estradiol to the brain for treating estrogen-responsive disorders. Sci Transl Med. 2015;7(297):297ra113. doi:10.1126/scitranslmed.aab1290

48. Cavaliere G, Trinchese G, Penna E, et al. High-Fat Diet Induces Neuroinflammation and Mitochondrial Impairment in Mice Cerebral Cortex and Synaptic Fraction. Front Cell Neurosci. 2019;13:509. doi:10.3389/fncel.2019.00509

49. Butler MJ, Perrini AA, Eckel LA. Estradiol treatment attenuates high fat diet-induced microgliosis in ovariectomized rats. Horm Behav. 2020;120:104675. doi:10.1016/j.yhbeh.2020.104675

50. Dash UC, Bhol NK, Swain SK, et al. Oxidative stress and inflammation in the pathogenesis of neurological disorders: Mechanisms and implications. Acta Pharm Sin B. 2025;15(1):15–34. doi:10.1016/j.apsb.2024.10.004

51. Zang X, Chen S, Zhu J, Ma J, Zhai Y. The Emerging Role of Central and Peripheral Immune Systems in Neurodegenerative Diseases. Front Aging Neurosci. 2022;14. doi:10.3389/fnagi.2022.872134

52. MacLusky NJ, McEwen BS. Oestrogen modulates progestin receptor concentrations in some rat brain regions but not in others. Nature. 1978;274(5668):276–278. doi:10.1038/274276a0

53. Sanathara NM, Moreas J, Mahavongtrakul M, Sinchak K. Estradiol upregulates progesterone receptor and orphanin FQ colocalization in arcuate nucleus neurons and opioid receptor-like receptor-1 expression in proopiomelanocortin neurons that project to the medial preoptic nucleus in the female rat. Neuroendocrinology. 2014;100(0):103–118. doi:10.1159/000363324

54. Li Z, Wei H, Li S, Wu P, Mao X. The Role of Progesterone Receptors in Breast Cancer. Drug Des Devel Ther. 2022;16:305–314. doi:10.2147/DDDT.S336643

55. Blaustein JD, Wade GN. Ovarian influences on the meal patterns of female rats. Physiol Behav. 1976;17(2):201–208. doi:10.1016/0031-9384(76)90064-0

56. Santollo J, Torregrossa AM, Eckel LA. Estradiol acts in the medial preoptic area, arcuate nucleus, and dorsal raphe nucleus to reduce food intake in ovariectomized rats. Horm Behav. 2011;60(1):86–93. doi:10.1016/j.yhbeh.2011.03.009

57. Baskin DG, Breininger JF, Bonigut S, Miller MA. Leptin binding in the arcuate nucleus is increased during fasting. Brain Res. 1999;828(1-2):154–158. doi:10.1016/s0006-8993(99)01252-4

58. Aponte Y, Atasoy D, Sternson SM. AGRP neurons are sufficient to orchestrate feeding behavior rapidly and without training. Nat Neurosci. 2011;14(3):351–355. doi:10.1038/nn.2739

59. Maejima Y, Kohno D, Iwasaki Y, Yada T. Insulin suppresses ghrelin-induced calcium signaling in neuropeptide Y neurons of the hypothalamic arcuate nucleus. Aging. 2011;3(11):1092–1097. doi:10.18632/aging.100400

60. Sims NA, Dupont S, Krust A, et al. Deletion of estrogen receptors reveals a regulatory role for estrogen receptors-β in bone remodeling in females but not in males. Bone. 2002;30(1):18–25. doi:10.1016/S8756-3282(01)00643-3

61. Davis KE, D. Neinast M, Sun K, et al. The sexually dimorphic role of adipose and adipocyte estrogen receptors in modulating adipose tissue expansion, inflammation, and fibrosis. Mol Metab. 2013;2(3):227–242. doi:10.1016/j.molmet.2013.05.006

62. Della Torre S, Mitro N, Fontana R, et al. An Essential Role for Liver ERα in Coupling Hepatic Metabolism to the Reproductive Cycle. Cell Rep. 2016;15(2):360–371. doi:10.1016/j.celrep.2016.03.019

63. Qiu S, Vazquez JT, Boulger E, et al. Hepatic estrogen receptor α is critical for regulation of gluconeogenesis and lipid metabolism in males. Sci Rep. 2017;7(1):1661. doi:10.1038/s41598-017-01937-4

